# A synthetic biology approach to bacterial transcription initiation: RNA aptamer based *in vitro* transcription assay for rapidly testing bacterial RNA polymerases, promoters and inhibitors

**DOI:** 10.64898/2026.08.11.744185

**Authors:** Tina Lanzmaier, Elena Reiterer, Melanie Merl, Andonita Ajdari, Karin Bischof, Günther Koraimann

## Abstract

We present a robust and versatile *in vitro* transcription (IVT) assay based on an optimized Broccoli RNA aptamer sequence. When paired with the fluorophore DFHBI-1T, this system enables real-time monitoring of multi-round transcription over several hours. To facilitate streamlined promoter analysis, we developed the pIVT3 plasmid backbone. The system was validated using both the single-subunit T7 RNA polymerase and the multi-subunit *Escherichia coli* RNA polymerase; notably, the activity of the *E. coli* enzyme remained strictly dependent on the presence of a σ factor and a cognate promoter. To optimize the signal-to-noise ratio, we incorporated two *rrnB*T1 terminators upstream of the promoter of interest. This modification effectively eliminated background transcription for weak promoters (P*livJ*) and prevented interference from read-through transcription in strong synthetic promoters (P*trc\**). Furthermore, we demonstrated the assay’s utility for drug discovery by characterizing the time- and dose-dependent inhibitory kinetics of rifampicin. Collectively, these results establish the Broccoli-based IVT system as a highly adaptable platform for quantifying promoter strength and screening small-molecule inhibitors of bacterial transcription.

**Graphical Abstract:** 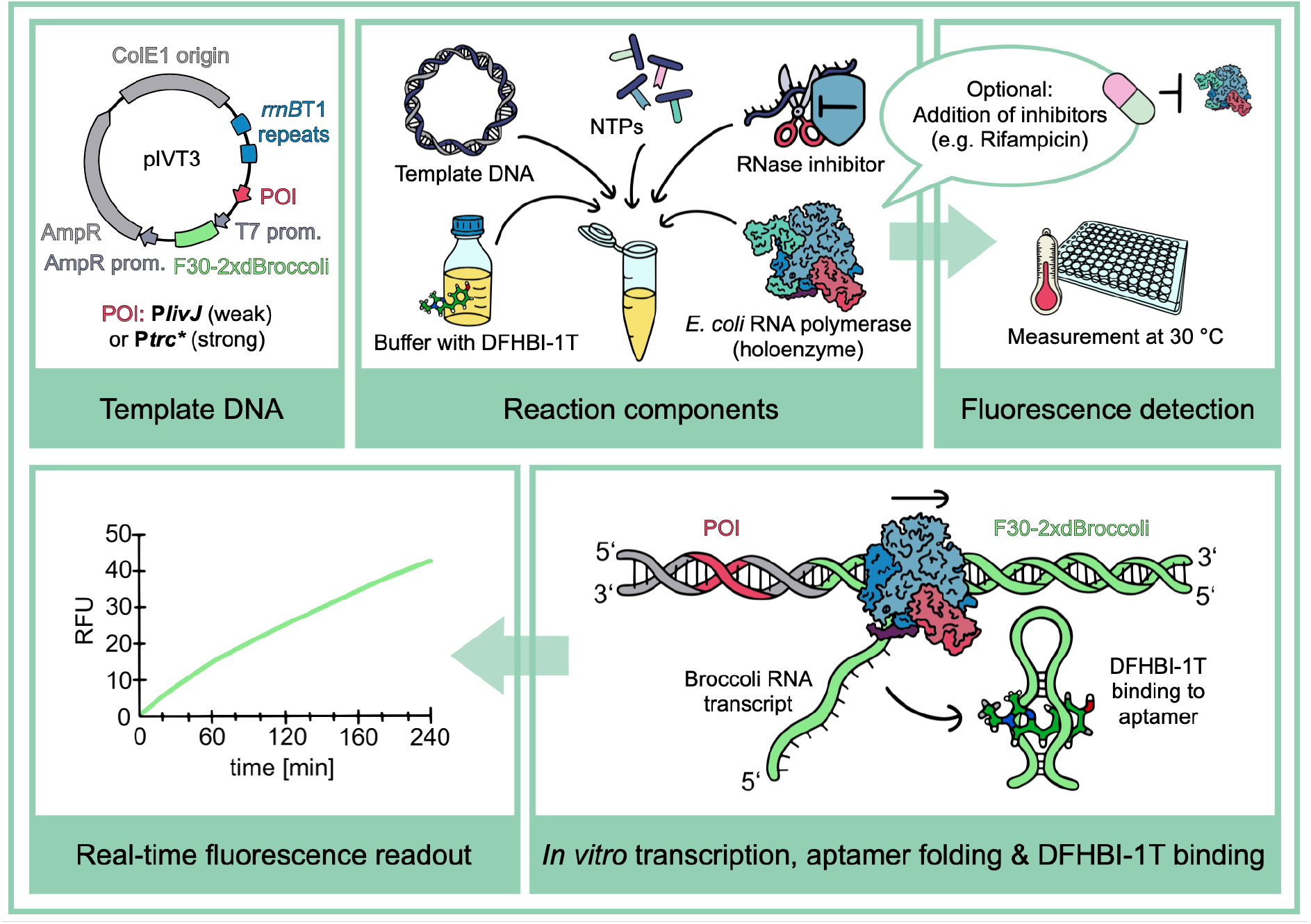

## Introduction

Bacterial transcription is a fundamental step in gene expression, executed by the multi-subunit enzyme RNA polymerase (RNAP). In bacteria, the core enzyme is capable of polymerizing single-stranded RNA from a double-stranded DNA template in the presence of nucleotide triphosphates (NTPs). In the model organism *Escherichia coli*, this core complex (E) consists of five subunits (2 α, β, β’, and ω) and possesses a molecular mass of approximately 400 kDa. To initiate transcription at specific DNA regions termed promoters, the core must associate with a σ factor (e.g., the housekeeping σ70 in *E. coli*). The interplay between various σ factors, transcriptional activators, and repressors governs global gene expression and metabolic adaptation in bacteria. For a recent review of the structural and mechanistic aspects of bacterial transcription initiation, see [1]. Initiation begins with the formation of a nucleoprotein complex, where the σ factor recognizes promoter elements to transition from a closed complex to an open complex (RPo). In RPo, the DNA strands are separated at the −10 element [2,3], exposing the template strand within the RNAP active site. Following initiation, the σ factor dissociates, and the core RNAP elongates the RNA transcript until it encounters a transcription terminator. Upon termination, the complex dissociates, releasing the nascent RNA, the DNA template, and the core RNAP, which then becomes available to re-associate with a σ factor [4]. Recent advances in Cryo-EM have provided high-resolution snapshots of these stages, including the *E. coli* holoenzyme-DNA open complex [5], coupled transcription-translation complexes [6], and pre-termination complexes [7].

Despite these structural insights, studying transcription initiation *in vitro*—the rate-limiting step of the cycle—remains challenging due to a lack of accessible and reproducible quantitative methods. Historically, *in vitro* transcription (IVT) assays relied on radioactive precursors, such as α-P^32^-ATP, requiring labor-intensive gel electrophoresis and autoradiography. To address these limitations, fluorescent light-up aptamers (FLAPs) have emerged as powerful tools for direct, real-time monitoring of transcription [8]. These RNA structural elements emit fluorescence upon binding a specific fluorophore, allowing for continuous data acquisition using standard fluorescence readers or real-time PCR machines. Building on this technology, we describe an IVT system utilizing the Broccoli RNA aptamer [9,10] integrated into optimized promoter test plasmids. By incorporating specific terminator elements to reduce background and read-through transcription, we demonstrate the system’s versatility through multi-round transcription studies of both strong and weak promoters. Furthermore, we validate the assay as a potential screening platform for RNAP inhibitors by characterizing the inhibitory effects of rifampicin during the early stages of transcription initiation.

## Materials and Methods

### Construction of pIVT3 plasmids

pIVT3 test plasmids (Fig. 3A) were designed based on ColE1 derived high copy number replicon and a selectable marker gene conferring resistance to ampicillin from the general purpose cloning vector pUC119 [11], two *rrnB*T1 transcriptional terminator elements from the promoter test vector pRS414 [12] to block readthrough from upstream transcription events, the POI (promoter of interest), and the DNA element encoding the Broccoli RNA aptamer [9]. Specifically, within this work, the 234 nt long F30-2xdBroccoli sequence was used, which consists of two dBroccoli sequences embedded in a F30 scaffold providing a very high and stable fluorescent signal in the presence of the fluorophore DFHBI-1T [9,10]. The Broccoli element was extended by the addition of an upstream T7 RNA polymerase promoter and amplified from plasmid pET28c-F30-2xdBroccoli - Addgene plasmid #66843 - [13]. PCR fragments representing these sequence elements and promoters of interest were combined by using NEBuilder HiFi DNA Assembly (NEB) according to the manufacturer’s instructions. A list of pIVT3 plasmids used in this study and inserted promoter sequences is given in the supplement (Table S1, Table S2). The DNA sequence of pIVT3ptrc* was verified by Oxford Nanopore Sequencing and is available in SnapGene (with annotated features) and Fasta formats in the figshare repository https://doi.org/10.6084/m9.figshare.33205119. This plasmid is available as DNA upon request and can be used to construct pIVT3 plasmids with other POIs cloned into the backbone by Gibson assembly [14].

### Plasmid DNA isolation

Plasmid DNA for IVT assays was purified from *E. coli* TOP10 cultures utilizing NucleoBond® PC 100 kit (Macherey-Nagel) according to the manufacturer’s instructions. In the final step the DNA pellet was solubilized in 100 µL purified water (Aqua bidest “Fresenius”, Fresenius Kabi, Austria). Concentrations of plasmid DNA were determined by Nanodrop ND-1000 measurements and were typically in the range between 200 ng/µL and 500 ng/µL.

### Proteins used in this study

RNA polymerases used in this study were either purchased from NEB (*E. coli* core enzyme, *E. coli* holoenzyme, bacteriophage T7 RNA polymerase) or self purified (*E. coli* core, *E. coli* σ70). For expression and purification of E RNAP we used the expression and purification system with the expression plasmid pIA900 [15]. Briefly, *E. coli* BL21(DE3) cells were grown in autoinduction medium at 16°C until an OD_600_ of 1 (± 0.1) was reached. After cell harvest by centrifugation cells were placed on ice for 30 min and then disrupted by sonification in lysis buffer (50 mM Tris-HCl, 500 mM NaCl, 1x cOmplete (Roche) protease inhibitor cocktail, 20 mM imidazole, 5% glycerol, 1 mg/mL lysozyme, pH 6.9). The cleared lysate was loaded onto a HisGraviTrap Talon (Cytiva) column, washed with binding buffer (50 mM Tris-HCl, 500 mM NaCl, 20 mM imidazole, 5% glycerol, pH 6.9). After elution with the same buffer containing 250 mM imidazole, fractions with the highest protein concentrations were pooled and dialyzed using Spectra-Por Float-A-Lyzer G2 (MWCO 8-10 kDa) against HiTrap Heparin A buffer (50 mM Tris-HCl, 75 mM NaCl, 0.5 mM EDTA, 5% glycerol, 1mM DTT, pH 6.9). The sample was subsequently injected into an ÄKTA FPLC system with a 1 mL HiTrap Heparin HP (Cytiva) column for a second affinity purification step. RNA polymerase core was eluted by applying a linear gradient from zero to 100 % HiTrap Heparin B buffer (same as A but with 1500 mM NaCl), the main peak appeared between 40 and 55 % buffer B. Peak fractions were pooled, the sample was concentrated with a buffer exchange to RNA polymerase reaction buffer (see below) by centrifugation through Amicon Ultra 10K Centrifugal Filter Units (Millipore). For storage at −20°C glycerol (87%) was added in a 1:1 ratio (v/v). For purification of σ70, *E. coli* BL21 (DE3) cells expressing N-terminally His6-tagged σ70 from pLNH12His6-σ70 [16] were used. Expression and purification was performed essentially as described [17] with the following modifications: Bacteria were cultured in 100 mL LB medium supplemented with 0.2% glucose and 200 µg/mL ampicillin. At OD_600_=0.7 protein expression was induced with 1 mM IPTG for 2.5 hours at 37°C. Cells were harvested by centrifugation and resuspended in 6 mL lysis buffer A (40 mM Tris-HCl, 300 mM KCl, 10 mM EDTA, 1x cOmplete (Roche) protease inhibitor cocktail, 0.2% sodium deoxycholate, pH 7.9) followed by sonification. Lysates were centrifuged for 30 min at 17600xg. Pellets with inclusion bodies were resuspended in lysis buffer B (same as A but containing 0.2% n-octyl-β-D-glucopyranoside as detergent), followed by sonification and centrifugation. This washing step was repeated with lysis buffer C (same as B but additionally containing 1 mM DTT). Finally, washed inclusion bodies were solubilized in denaturing buffer (40 mM Tris-HCl, 6 M guanidinium-HCl, 10 mM MgCl_2_, 1 mM EDTA, 2 mM DTT, 10% glycerol, pH 7.9). After incubation on ice for 30 min and a centrifugation step, the supernatant was adjusted to 500 mM NaCl and 5 mM imidazole and then loaded onto a 1 mL HisTrap HP column (Cytiva). Bound proteins were washed (50 mM Tris-HCl, 500 mM NaCl, 5 mM imidazole, 6 M guanidine-HCl, pH 7.9) and subsequently eluted with the same buffer containing 500 mM imidazole. Fractions containing protein with renatured His6-σ70 were pooled and PMSF was added (1 mM final concentration). Protein refolding was performed by dialysis against renaturation buffer (50 mM Tris-HCl, 200 mM KCl, 10 mM MgCl_2_, 10 µM ZnCl_2_, 1 mM EDTA, 1 mM DTT, 30% glycerol, pH 7.9). After increasing the final glycerol concentration to 45%, aliquots of the samples were stored at −20°C and used for reconstitution of RNAP holoenzyme.

Protein quality control was performed by SDS-PAGE using precast NuPAGE 4-12% Bis-Tris protein gels (Invitrogen) and subsequent Coomassie staining using Colloidal Blue Staining Kit (Invitrogen). Protein concentrations were determined by the Bio-Rad Protein Assay (Bio-Rad) according to the method developed by Bradford [18]. An SDS-PAGE with purified proteins and protein complexes, alongside purchased RNA Polymerases is shown in the supplement (Figure S1).

### Standard IVT assays with self-made RNA polymerase and reconstituted Eσ70

#### Reaction conditions

IVT assays were performed in a volume of 20 µL containing 15 nM plasmid DNA, 30 nM *E. coli* RNAP supplemented with a 2.5-fold surplus of σ70, RNA polymerase reaction buffer (“RNAP Buffer” according to a NEB recipe: 40 mM Tris-HCl, 150 mM KCl, 10 mM MgCl_2_, 1 mM DTT, 0.01% Triton® X-100, pH 7.5), 12.5 µM DFHBI-1T (Sigma-Aldrich), 1U/µL RNase Inhibitor, Murine (NEB), and 0.5 mM of each NTPs (Thermo Fisher Scientific).

#### Reaction set up and real-time measurement

RNAP holoenzyme (150 nM) was reconstituted from purified core and σ70 (375 nM) in reconstitution buffer (50 mM Tris-HCl, 200 mM KCl, 10 mM MgCl_2_, 1 mM DTT, 1 mM EDTA, 0.001 mM ZnCl_2_, 10% Glycerol, pH 7.9). After incubation for 15 min at 30°C, 4 µL of this mix was combined with 4 µL 5x RNAP Buffer containing 62.5 µM DFHBI-1T, 0.5 µL RNase Inhibitor (40 U/µL) and 4.5 µL purified water (Aqua bidest “Fresenius”, Fresenius Kabi, Austria) yielding “RNAP Master Mix”. 13 µL of this RNAP Master Mix was then combined with 5 µL 60 nM plasmid DNA (in purified water) and incubated for 15 min at RT in the dark to allow RPo (RNA polymerase open complex) formation. To allow transcription to initiate and proceed, 2 µL of a NTP mix (5 mM of each ATP, CTP, GTP, UTP) was added resulting in a final volume of 20 µL. All steps were performed on ice or a cold block unless otherwise indicated. All pipetting steps were performed using RNase-free pipette tips with filters (Biozym) to avoid contamination with RNases. Mixing of components was done by gently pipetting up and down for 3-4 times. Concomitant with the final step, IVT reactions were transferred to 0.1 mL PCR tubes for Rotor-Gene real-time PCR detection systems. For recording transcription and Broccoli RNA production in real-time we utilized a RotorGene RG-6000 machine (Corbett Research) with the tubes placed in a rotor with 72 positions. Fluorescence was recorded in the green channel (excitation 470 nm, emission 510 nm) during the whole run once per cycle. One cycle had a duration of 1 min 23 sec. The temperature was kept at 30°C. After the run (typically 210 cycles) samples were transferred to −20°C until further use.

### Prototype IVT assays with purchased RNA polymerases

Prototype assays for validation of the IVT system contained 15 nM plasmid DNA and 75 nM T7 RNA polymerase (NEB) for assaying transcription from the T7 promoter present on the pIVT plasmids. Alternatively, E (“core”) or Eσ70 (“holo”) purchased from NEB were used at final concentrations of 50 or 60 nM, respectively. 1 µL of purchased RNAP was mixed with 4 µL 5x RNAP Buffer containing 62.5 µM DFHBI-1T, 1 µL RNase Inhibitor (40U/µL) yielding “RNAP Master Mix”. 6 µL of this mix was combined with 12 µL plasmid DNA to allow RPo formation, samples were further processed as described above.

### Data processing and statistics

Recordings from the IVT run were exported from the Rotor-Gene Q Series Software 2.3.1 in CSV format and further processed in Google sheets. To account for the observed differences in reaction kinetics, four “time windows” were defined which were then used to calculate slopes reflecting transcription initiation (or promoter activity) according to the following formula:

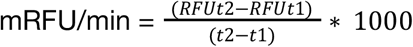

RFU: relative fluorescence units

t: time after initiation of transcription in min

The resulting values (mRFU/min) were used to compare different sample conditions. Statistical analyses and visualizations were performed using numiqo online statistics calculator (https://numiqo.com). Data analyses and statistics for results of experiments shown in Figs. 2 and 4 are detailed in the supplement (Data S1 and Data S2), respectively.

### DNA/RNA agarose gel electrophoresis

RNA reaction products and input plasmid DNA can be analysed after the real-time measurements by DNA/RNA agarose gel electrophoresis. Typically, 10 µL from the IVT reactions were mixed with DNA loading buffer and run on a standard 1% non-denaturing agarose gel alongside with suitable DNA molecular weight markers. First, RNA containing the Broccoli aptamer was visualized by staining the gel with a buffer (40 mM Tris-HCl pH 7.4, 100 mM KCl, 1 mM MgCl_2_) containing 10 µM DFHBI-1T at room temperature for 1 h. After documentation with a ChemiDoc MP system (BioRad) with blue epi-illumination (488 nm LED) together with a 530/28 filter the gel was stained again with SybrSafe (1:10.000 in 0.5x TAE buffer, 1 h at room temperature) to allow detection of total RNA and DNA. Images of the gels were again generated by using the ChemiDoc MP system with the same settings as above.

## Results

### Multi-round transcription by T7 or *E. coli* RNAP can be monitored quantitatively in real-time and qualitatively by analysis of reaction products

pIVT plasmids as shown in Figure 3A were directly used as substrates in an IVT experiment together with an active RNAP and NTPs in a buffer containing DFHBI-1T. Transcriptional activity was monitored during IVT reactions in real time using a RT PCR machine over a period of several hours (Fig. 1A), which allowed us to capture dynamic aspects of the IVT reactions [8]. In a first series of experiments with pIVT plasmids containing the strong synthetic promoter designated P*trc*\*, we tested the properties and robustness of the system. The P*trc*\* promoter was derived from the *livJ* promoter (−86 to +32 from *E. coli*) by replacement of the sequence from −35 to −10 with the *trc* promoter core sequence element while leaving the original *livJ* flanking sequence context, except for a 1 nt deletion upstream of −35 (see Figure 3B and table S1). Thus, the promoter designated P*trc\** in this work is a hybrid of *livJ* UP and DOWN elements combined with the core of the original *trc* promoter [19]. The *trc* promoter is a 17 bp spacer derivative of the *tac* promoter which by itself is a hybrid of *lacUV5* and *trp* promoter sequences from *E. coli* [20].

**Fig. 1:**
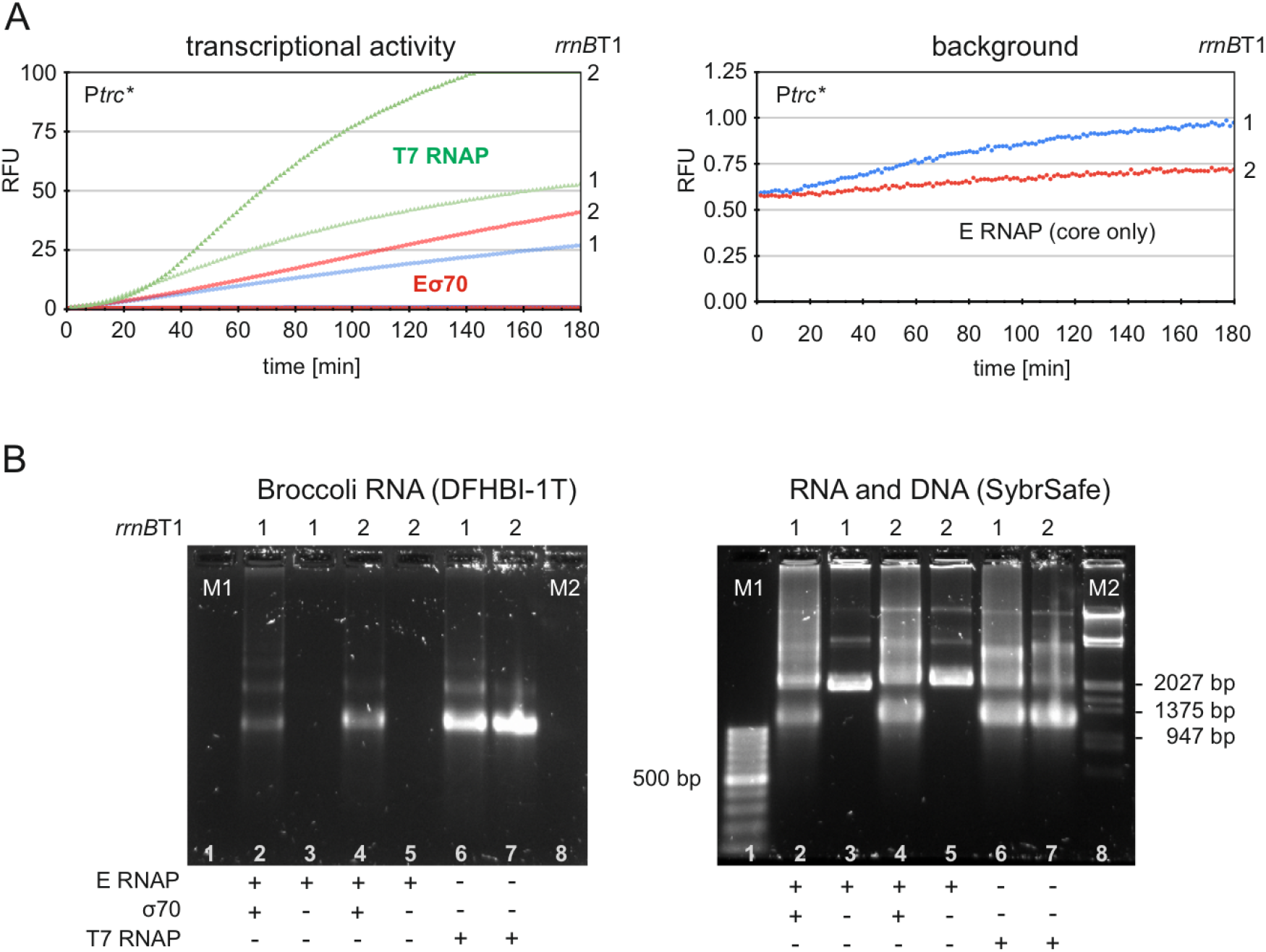
Validation of the IVT system and its readouts. RNA Polymerases purchased from NEB were used. **A:** Real time recordings of fluorescence signals generated by transcription events. Transcription from the strong P*trc*\* or T7 promoters by Eσ70 (60 nM) or T7 RNAP (75 nM), respectively, can be readily measured by a real-time PCR machine at a constant temperature (usually 30°C) over a time period of 180 minutes. The slope of the curve is proportional to the promoter strength because initiation is the limiting step in transcription. Two *rrnB*T1 terminators can efficiently reduce read-through transcription and block interference with initiation events at the promoters. The panel to the right shows the very low background activity produced by E core RNAP (50 nM, no sigma factor). **B:** Post reaction analysis by agarose gel electrophoresis. The products (RNA molecules containing the Broccoli aptamer) of the IVT reactions are visualized by staining with DFHBI-1T (left panel) and fluorescence detection. Fluorescent RNA is only produced by Eσ70 (lanes 2 and 4) or T7 RNAP (lanes 6 and 7) but not by E RNAP (lanes 3 and 5) in the IVT reaction. Two *rrnB*T1 terminators can efficiently reduce read-through transcription (e.g.: compare lanes 2 and 4). M1 and M2 (lanes 1 and 8) contain DNA standards which are only visible after SybrSafe staining (right panel) of the same agarose gel. In that case, both RNA and the DNA are stained and visualized. The input plasmid DNA is best seen in the lanes with E RNAP only. Some sizes of marker DNA fragments (in base pairs, bp) are indicated for lanes M1 and M2, respectively.

In a first series of experiments we utilized RNA polymerases purchased from NEB. We used single subunit RNAP from bacteriophage T7 (T7 RNAP) and compared it to multi-subunit RNAP from *E. coli* (Eσ70 or E) to check for general functioning of the IVT system. Reaction conditions were as described in the Materials and Methods section. Both T7 RNAP and Eσ70 produced transcripts which could be readily monitored in real time over a time period of at least three hours (Fig. 1A). To verify the generation of RNA, reaction products were subsequently analyzed by non-denaturing agarose gel electrophoresis. Sequential staining of the gel, first with a DFHBI-1T solution and afterwards with the DNA/RNA specific SybrSafe stain revealed the presence of RNA transcripts containing the Broccoli FLAP alone or together with the input plasmid DNA, respectively (Fig. 1B). In case of the plasmid constructs with 2 *rrnB*T1 terminators, most transcripts had a length of approximately 2500 nt and both T7 RNAP and Eσ70 appeared to efficiently terminate at the *rrnB*T1 elements. Furthermore, E-RNAP (core only) did not produce measurable transcripts, confirming the notion that transcription by a bacterial multi-subunit RNAP from a circular DNA molecule is strictly dependent on a promoter and a cognate sigma factor (in this case σ70).

Interestingly, in case of the strong T7 and P*trc*\* promoters, the additional second *rrnB*T1 terminator resulted in even higher final RNA amounts and higher overall transcription activities. We attribute this effect to reduced interference by transcriptional read-through which has been shown to negatively affect efficient re-initiation [21]. To further explore this possibility we subsequently investigated the effects of the absence or presence of one or two *rrnB*T1 elements using a strong or a weak promoter for comparison.

### Two *rrnB*T1 terminators provide sufficient protection from upstream transcription and allow to monitor transcription from the weak *livJ* promoter

To investigate the effect of the presence of *rrnB*T1 terminators in case of a weak promoter we used P*livJ* which is known as an Eσ70 dependent promoter [22]. It drives transcription of the *livJ* gene encoding a periplasmic binding protein of an ABC transporter system specific for branched-chain amino acids and phenylalanine [23,24]. Transcription of *livJ* in its chromosomal context can be repressed by leucine-responsive regulatory protein (Lrp) in the presence of branched-chain amino acids [25]. On the other hand, transcription of *livJ* in *E. coli* can be activated by nutrient limitation (stringent response) via the secondary channel RNAP binding protein DksA together with the alarmone ppGpp [26,27] or by the F-like plasmid-encoded transcriptional modulator TraR [28]. It is not clear, however, whether this activation is a direct effect or an indirect effect due to fewer Eσ70 engaged in transcription from promoters that are negatively affected by the stringent response [27–29]. Both P*livJ* and P*trc*\* promoters were then used together with the pIVT backbone containing none, one or two *rrnBT1* terminators, respectively (Fig. 2). Transcriptional activity of the highly active P*trc*\* promoter is significantly negatively affected only at later time points if no terminator is present upstream of the promoter (Fig. 2D). There is approximately a 1.5-fold reduction in transcriptional activity from the first time window (20-80 min) to the last one (200-260 min) which is highly significant (p < 0.001). Accordingly, the total amount of Broccoli-containing RNA is highest with two *rrnB*T1 terminators (Figs. 2B&C). With two *rrnB*T1 terminators preceding the promoter sequence the negative interference effect is virtually eliminated and only one major RNA species with a length of approximately 2400 to 2600 nt is visible as a prominent band in the agarose gel after staining with DFHBI-1T (Fig. 2C). This IVT phenotype is consistent with the interpretation that most transcription events start at P*trc*\* and terminate at one of the *rrnB*T1 terminators. Notably, during the whole duration of the fluorescence recording (240 min) there was no significant decrease in the activity of P*trc*\* as expressed in mRFU/min showing that the conditions used were not limiting in the multi-round transcription assay (Fig. 2D).

**Fig. 2:**
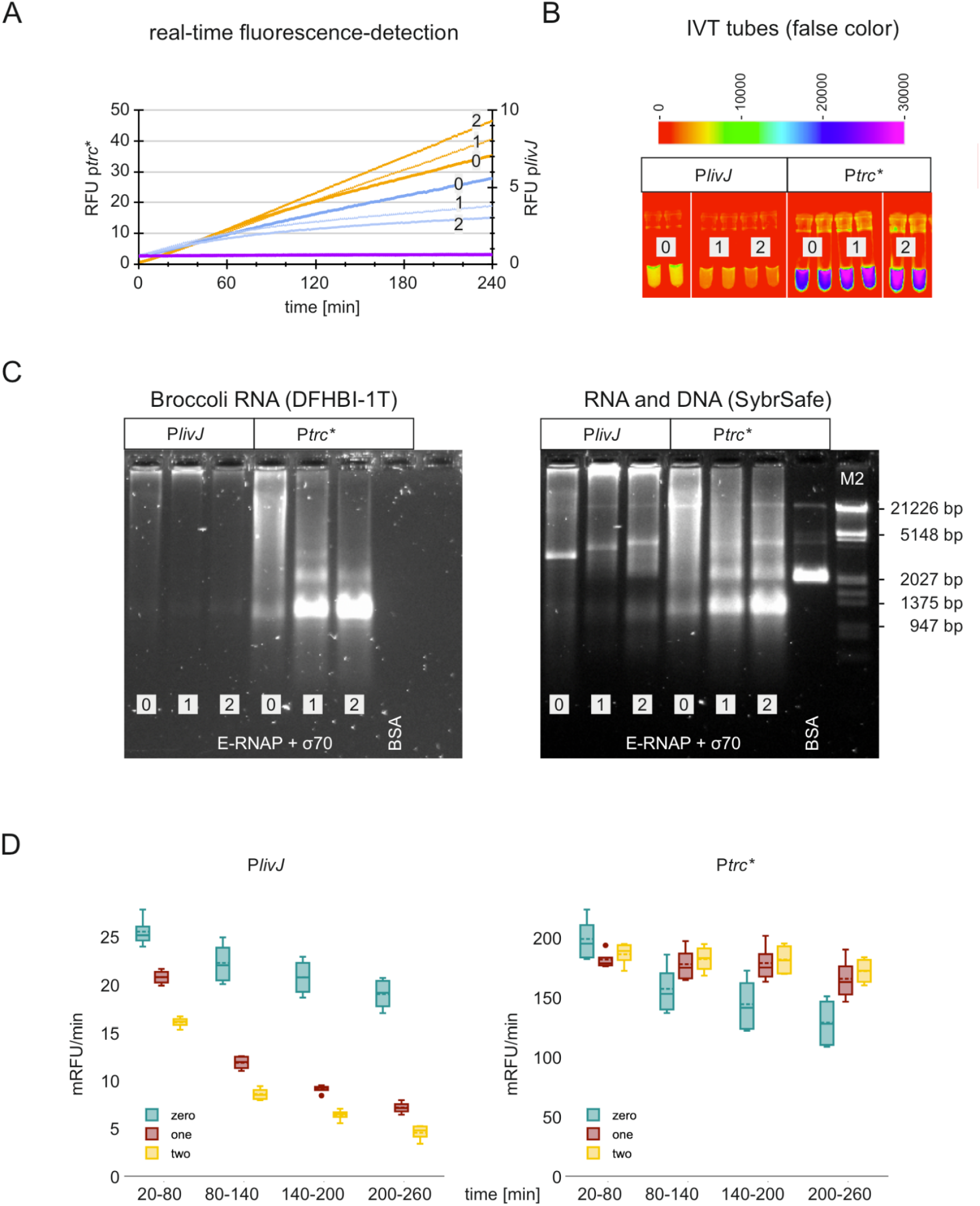
Opposing effects of the presence of two rrnBT1 terminator sequences on overall transcription rates from P*livJ* and P*trc*\* promoters. **A:** Real-time fluorescence-detection of DFHBI-1T-bound Broccoli aptamer RNA from IVT reactions employing promoters P*livJ* (blue) or P*trc\** (orange). The violet line represents the reaction with BSA instead of Eσ70. **B:** Fluorescence image obtained from IVT tubes after the incubation step **C:** Agarose gel electrophoresis as described in detail in figure 1. **D:** Quantitative analysis and comparison of transcription rates (mRFU/min) determined for different periods of time after the start of IVT reactions. For a detailed description and interpretation see the main text. The absence (“0”, “zero”) or presence of one (“1”, “one”) or two (“2”, “two”) *rrnB*T1 terminators is indicated. Data shown in **D**: are the result of two independently carried out IVT experiments with duplicate samples in each run. Data used to create the box plot, analyses and statistics are presented in the supplement: Data S1 (Fig.2).

In contrast, the approximately 10-fold weaker *livJ* promoter displayed decreasing activity with prolonged incubation times, possibly indicating sensitivity towards changes in the DNA topology induced by transcribing RNA polymerase. This effect was particularly obvious in the presence of two *rrnB*T1 terminators indicating that transcripts originating from other promoters present on the template pIVT plasmid were efficiently terminated. This is also clearly visible in Fig. 3C because of the absence of longer RNAs in the upper part of the agarose gel. To further define background levels of Broccoli RNA signals arising from transcription events we cloned a DNA fragment without an obvious promoter element. Such a DNA fragment (VC16S, table S1) cloned into a pIVT3 plasmid (with 2 *rrnB*T1 terminators) resulted in a very low background signal at 260 min: 0.73 RFU ± 0.04; promoter activity (time window 20-80 min): 1.61 ± 0.41 mRFU/min (n=7), similar to what we observed in case of E-RNAP without the sigma factor (Fig. 1). This strongly suggests that the *rrnB*T1 elements in tandem can effectively shield transcription from upstream promoters located on the circular plasmid, e.g. the AmpR promoter. We infer from these results that signals above the values obtained for the VC16S “non-promoter” sequence solely arise from the promoter of interest (POI) cloned into the pIVT3 plasmid.

**Fig. 3:**
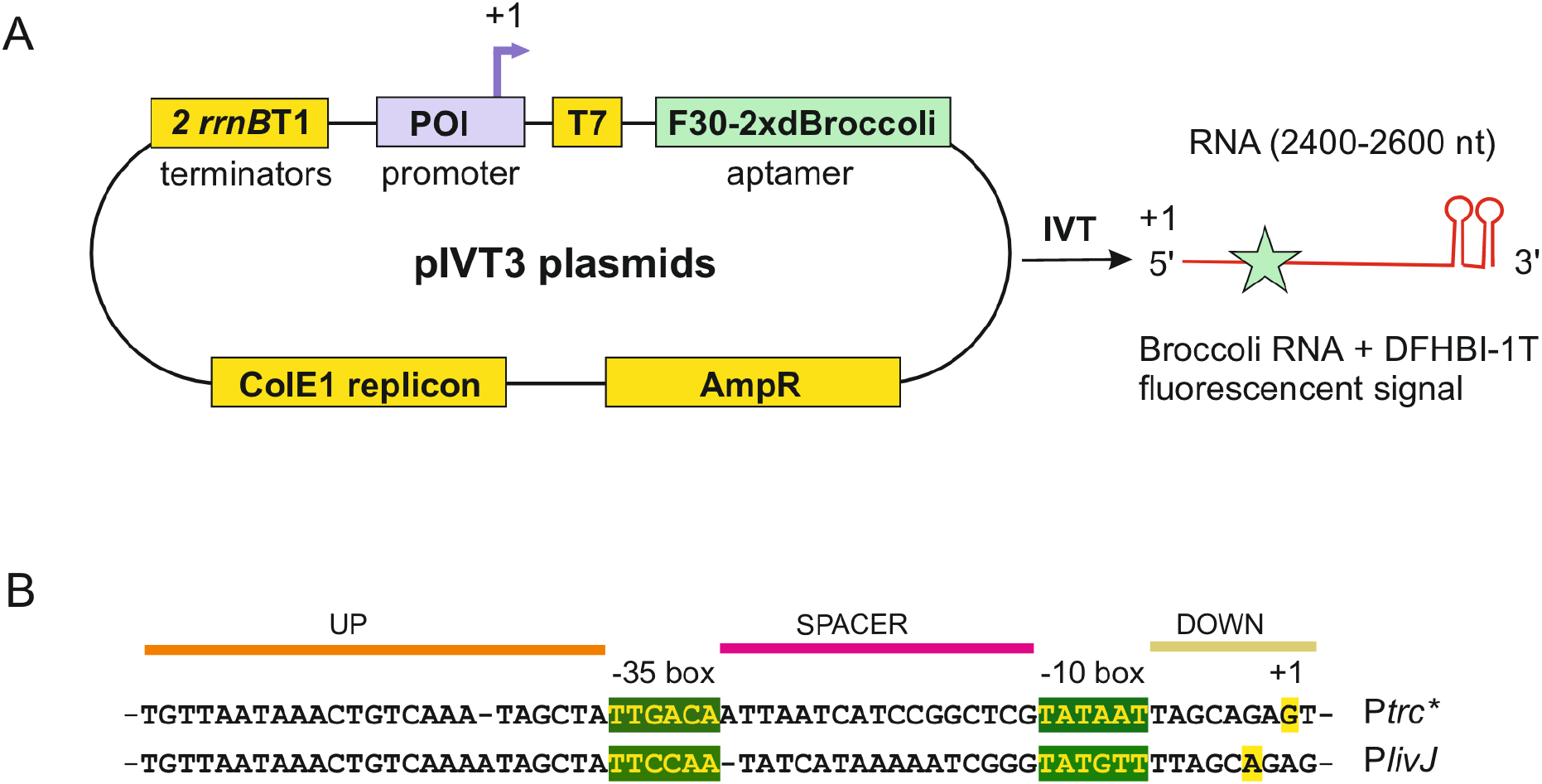
Plasmid architecture and promoter sequences. **A:** A circular plasmid DNA template is provided in order to minimize background transcription by E RNAP. Key components of the pIVT3 plasmid series are the POI (promoter of interest), a T7 promoter box for test purposes, a F30-2xdBroccoli element for the RNA aptamer, selectable resistance gene marker cassette (AmpR), a replicon for autonomous plasmid replication (ColE1) and finally two *rrnB*T1 terminators (2x*rrnB*T1). In the *in vitro* transcription reaction (IVT) RNA is produced containing the Broccoli RNA aptamer which emits a fluorescent signal upon binding of the fluorophore DFHBI-1T. **B:** DNA sequences of promoters used in this study. The sequences are aligned to their respective −35 and −10 regions (= core promoter, highlighted in green). UP/DOWN elements as well as the spacer sequences are shown. Due to its construction, P*trc\** has the same UP and DOWN elements as P*livJ* - except for a single base deletion. Transcription start sites (+1) are highlighted in yellow and were taken from the RegulonDB database [30] version 13.6.0 for P*livJ*, or were determined by 5’RACE of in vitro transcribed RNA for P*trc\** (data not shown). Plasmids and the POI DNA sequences inserted into the respective pIVT3 promoter test plasmid from this study are listed in the supplement (Table S1). The complete pIVT3Ptrc* plasmid sequence is available in the public figshare repository https://doi.org/10.6084/m9.figshare.33205119.

### Rifampicin can completely block transcription from the highly active *trc*\* promoter

In the experiments described above we established that pIVT3 plasmids as depicted in Fig. 3A are well suited to quantitatively and qualitatively monitor multiround transcription events that essentially reflect promoter activity. To establish that the system can also be used to screen for potential new inhibitors of bacterial RNAP we applied the well characterized inhibitor of transcription, rifampicin, at concentrations of 1 or 5 µM to our standard IVT reactions with components of *E. coli* RNAP and pIVT3 plasmid containing P*trc*\* as a promoter. These concentrations are well above the IC50 value of 15 nM determined recently (Jensen et al., 2023). The experimental setup was chosen to show that inhibition of transcription can be detected independently of the step in which rifampicin is added to the reaction. For that purpose, the drug was either added to the reconstitution mixture of core RNAP and sigma factor, the NTP mixture used to start the IVT reaction (after formation of Eσ70 and addition of plasmid template DNA), or after about two hours of ongoing transcription and fluorescence recording (Fig. 4). Transcription could be inhibited regardless of the step rifampicin was introduced into the system, albeit with quantitatively different outcomes. First of all, addition of rifampicin completely blocked transcription when added to the transcription buffer and Eσ70 in a step before the addition of DNA template (Fig. 4A). There is no detectable aptamer containing RNA and the measured transcriptional activity is below the background activity as determined above. Nevertheless, a part of the supercoiled plasmid DNA appears to be shifted to two distinct Eσ70-DNA complexes (which do not appear in the control lane without Eσ70) with one of them very likely representing the RPo form (open complex). This is consistent with the fact that rifampicin blocks the elongation step in transcription by occupying the RNA exit channel in the beta subunit [31]. Second, rifampicin, if added to the NTP mixture used to start the IVT reaction, only strongly inhibited transcription after a “lag” phase (Fig. 4B). The transcriptional activity in the time window 20-80 min was readily measurable but reduced in a rifampicin concentration dependent manner. We infer that the residual transcriptional activity represents a “single-round” transcriptional event where rifampicin could not block the first round of transcription from P*trc*\* after the formation of RPo. As a consequence, full-length transcripts can be seen in the post-reaction agarose gel electrophoresis in Fig. 4B as weak bands (1 or 5 µM rifampicin). In the third experimental setup transcription was allowed to proceed for about 2 hours before rifampicin was added to the samples. As can be seen in Fig. 4C, transcription rates dropped significantly and reached similar levels when compared to the experiment when rifampicin was added to the NTP mix. Taken together, our results confirm the notion that rifampicin can block transcription at an early initiation step of transcription. Thus, inhibition was most effective when rifampicin was added to Eσ70 before the addition of DNA. Data and statistical analyses for this set of experiments are provided in the supplement: Data S2 (Fig.4).

**Fig. 4:**
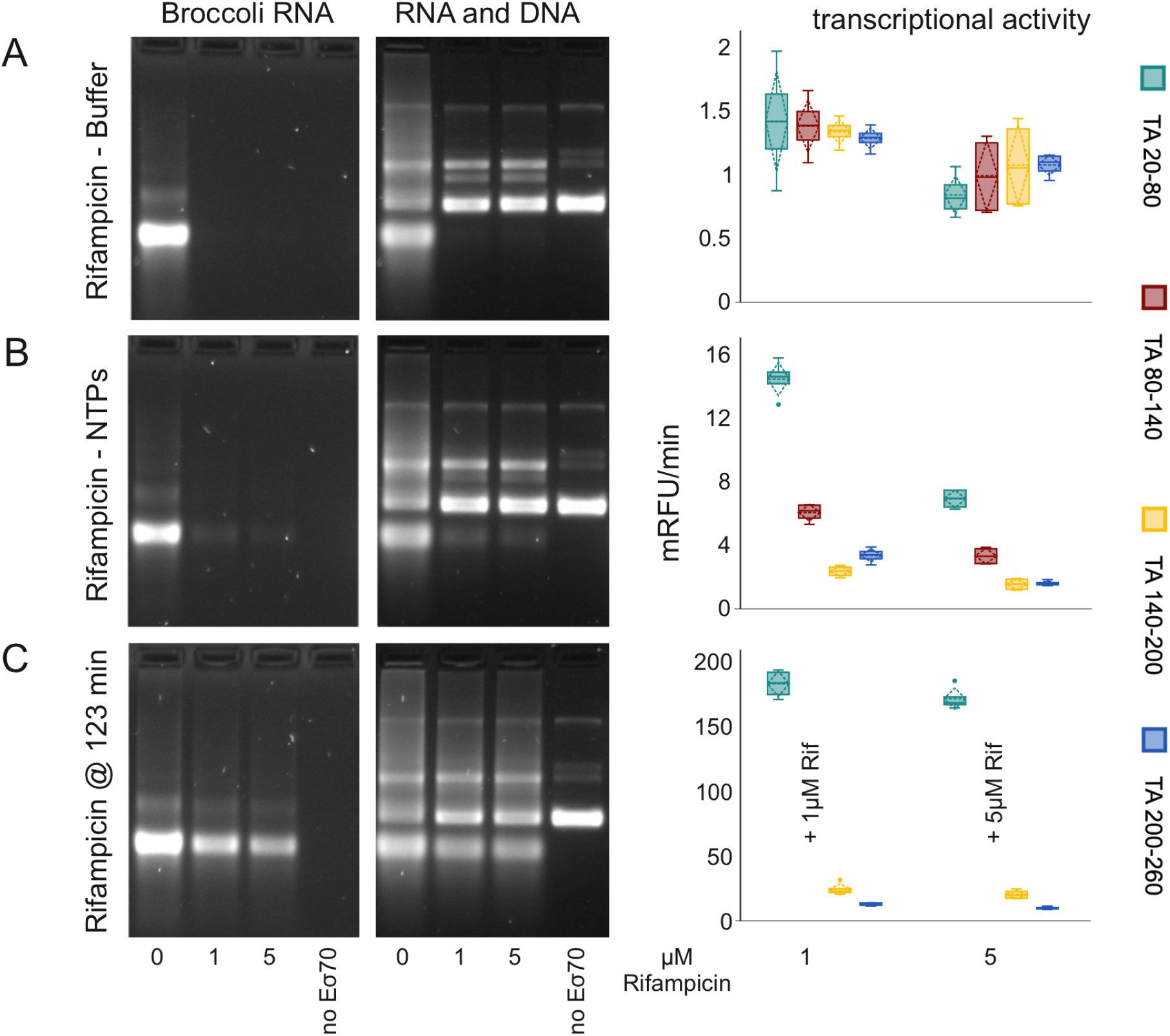
Inhibition of transcription by rifampicin at different stages of the IVT assay. Left: Post reaction agarose gel electrophoresis for visualization of Broccoli RNA (DFHBI-1T staining) and both RNA and DNA (SybrSafe staining). Lanes labelled with “0” (“zero”), “1” (“one”) or “5” (“five”) depict results from reactions without rifampicin, with 1 µM rifampicin or with 5 µM rifampicin, respectively. Control reactions without Eσ70 are indicated by “no Eσ70”. Right: Comparison of transcription rates (mRFU/min) determined for different periods of time after the start of IVT reactions. The concentration of rifampicin (1 µM or 5 µM) within the reaction mixtures is indicated. **A:** Agarose gel electrophoresis and transcriptional activity for samples to which rifampicin was added to transcription buffer and Eσ70 mixture prior to addition of template DNA. **B:** Results for IVT reactions with rifampicin added to the NTP mixture. **C:** Results for addition of Rifampicin after approximately two hours of ongoing transcription reaction. Data used to create the box plots, analyses and statistics are presented in the supplement: Data S2 (Fig.4).

## Discussion

The IVT system presented here provides a versatile framework for real-time, quantitative monitoring of bacterial transcription. Central to this approach is the pIVT plasmid backbone, which utilizes a tandem arrangement of *rrnB* T1 terminators to minimize background interference. For weak promoters, this shielding prevents non-specific upstream transcription from obscuring the results, while for highly active promoters like P*trc\**, it prevents transcriptional interference that could otherwise diminish activity over time. This architectural optimization ensures that the observed fluorescence signal accurately reflects the activity of the promoter of interest. Furthermore, by using supercoiled circular DNA as the template, we achieved a strict dependence on σ factor for initiation, more closely mimicking the requirements of native bacterial transcription compared to linear templates.

While the Broccoli RNA aptamer and similar fluorescent reporters have been utilized in previous IVT setups [8,32,33], our method offers several distinct practical advantages. The inclusion of dual upstream terminators ensures that the promoter of interest is isolated from the rest of the plasmid environment, preventing confounding signals from other elements like the ampicillin resistance promoter. Additionally, the high sensitivity of the system allows for the use of low concentrations of plasmid template (15 nM) and holoenzyme (30 nM), thereby significantly reducing reagent costs and maintaining stable reaction conditions over extended periods.

Taken together, our IVT system is robust and can easily be modified or scaled up to allow high-throughput screening for inhibitors of bacterial transcription by single- or multi-subunit RNA polymerases. It can also be used to test various promoters and promoter elements as well as the impact of different sigma factors, activators and inhibitors of transcription.

