## Supplemental information for "A synthetic biology approach to bacterial transcription initiation: RNA aptamer based *in vitro* transcription assay for rapidly testing bacterial RNA polymerases, promoters and inhibitors"

#### Table S1: Bacterial strains and plasmids

a) *Escherichia coli* strains

| strain designation | genotype | description / use |
|---|---|---|
| TOP10 | F–*mcr*A Δ(*mrr-hsd*RMS-*mcr*BC) φ80*lac*ZΔM15 Δ*lac*X74 *rec*A1 *ara*D139 Δ(*ara-leu*)7697 *gal*U *gal*K λ–*rps*L(StrR) *end*A1 *nup*G | transformation of *in vitro* generated plasmids; plasmid DNA preparation |
| BL21(DE3) | F–*omp*T *hsd*SB (rB–, mB–) *gal dcm* (DE3) | T7 RNA polymerase based system for protein expression and purification |

b) Recombinant plasmids from this study

| plasmid name | features / description | size (bp) |
|---|---|---|
| pIVT1PlivJ | $P_{livJ}$ - $P_{T7}$ *ampR ColE1-ori F30-2xdBroccoli* | 2421 |
| pIVT2PlivJ | $P_{livJ}$ - $P_{T7}$ *ampR ColE1-ori F30-2xdBroccoli rrnB*T1 (1) | 2671 |
| pIVT3PlivJ | $P_{livJ}$ - $P_{T7}$ *ampR ColE1-ori F30-2xdBroccoli rrnB*T1 (2) | 2854 |
| pIVT1Ptrc* | $P_{trc*}$ - $P_{T7}$ *ampR ColE1-ori F30-2xdBroccoli* | 2420 |
| pIVT2Ptrc* | $P_{trc*}$ - $P_{T7}$ *ampR ColE1-ori F30-2xdBroccoli rrnB*T1 (1) | 2670 |
| pIVT3Ptrc*[1] | $P_{trc*}$ - $P_{T7}$ *ampR ColE1-ori F30-2xdBroccoli rrnB*T1 (2) | 2853 |
| pIVT3VC16S | VC16S - $P_{T7}$ *ampR ColE1-ori F30-2xdBroccoli rrnB*T1 (2) | 2929 |

[1] The DNA sequence of plasmid pIVT3Ptrc* was verified using Oxford Nanopore sequencing. The data is accessible through the figshare repository: https://doi.org/10.6084/m9.figshare.33205119.

### Table S2: Promoter sequences and features. Sequences of DNA fragments (promoters of interest = POI) cloned into the pIVT promoter test plasmids are shown.

| name[1] | origin[2] | start[3] | stop[3] | length (bp) | DNA Sequence[4] |
|---|---|---|---|---|---|
| $P_{livJ}$ | *E. coli* MG1655 | -86 | +32 | 118 | ccggcaaaacgccaatccccacgcagattgttaataaac tgtcaaaatagcta**TTCCAAtatcataaaaatcgggTAT GTTttagcA**gagtatgctgctaaagcacgggtagtcatg c |
| $P_{trc^*}$ | *E. coli* composite/ synthetic | -88 | +29 | 117 | ccggcaaaacgccaatccccacgcagattgttaataaac tgtcaaatagcta**TTGACAattaatcatccggctcgTAT AATtagcagaG**tatgctgctaaagcacgggtagtcatgc |
| VC16S | *V. cholerae* | -160 | +33 | 193 | caatctgtgtgggcactcgttgatgataatcaaaaaaga tttatcaatgaactgagtgaccatttgaatgagcaatca ttcagcacagtcaattcactatcgaaagatagtatcagt attcattgagccgaagcgaaagcttcacaaaacttttaa ttgaagagtttgatcatggctcagattgaacgctggc |

1) name of the promoter element
2) origin of the promoter element indicates the genetic origin (organism) of the cloned DNA sequence (POI in pIVT3 plasmids)
3) start and stop positions relative to the transcription start site (+1) are given. In case of VC16S, the positions relative to mature 16S rRNA are indicated, no DNA sequence resembling a promoter element is present in this case
4) DNA sequence: Non-coding strand DNA sequence is shown; the core promoter sequence is shown in bold letters, core elements (-35, -10, +1) are in uppercase letters

### Figure S1: SDS PAGE of purified proteins and protein complexes

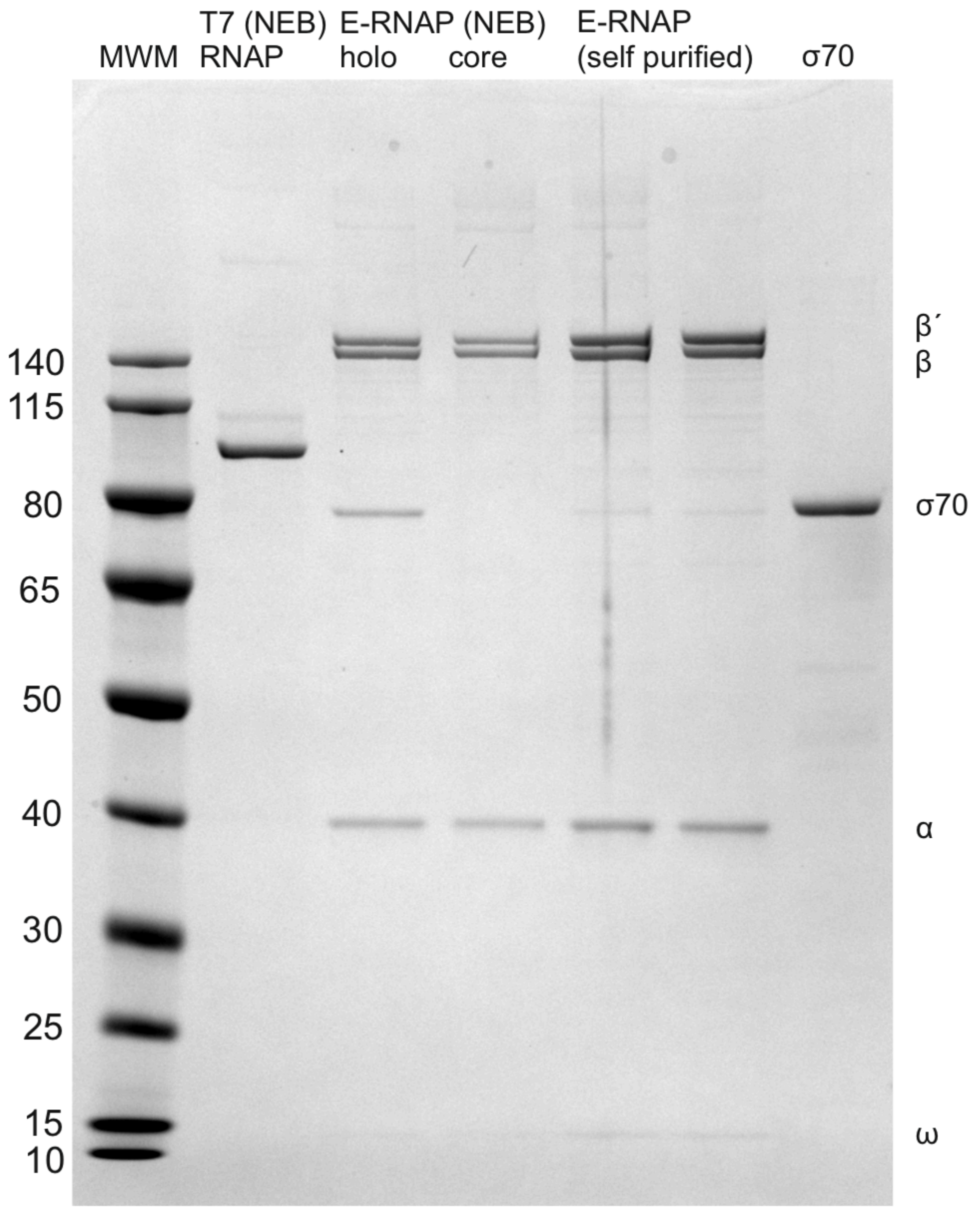


**Figure S1:** Coomassie stained SDS PAG (NuPAGE 4-12%, MOPS buffer) showing purified proteins and protein complexes used in the IVT experiments. **Lane 1**: Thermo Scientific PageRuler Prestained Protein Ladder, 10-180 kDa. Protein apparent molecular weights are indicated on the left according to Thermo Scientific Pub. No. MAN0011772 Rev. Date 19 June 2019 (Rev. C.00); **Lane 2**: Bacteriophage T7 RNA polymerase (NEB M0251); **Lane 3**: *E. coli* RNA polymerase, holoenzyme (NEB M0551); **Lane 4**: *E.coli* RNA polymerase, coreenzyme (NEB M0550); **Lanes 5,6**: Purified *E. coli* RNA polymerase as described in the Materials and Methods section, aliquots of two purification batches; **Lane 7**: Purified σ70 from *E. coli* as described in the Materials and Methods section. Subunits of *E. coli* RNA polymerase are indicated on the right side.

#### Data S1 (Fig.2):

**Opposing effects of the presence of two rrnBT1 terminator sequences on overall transcription rates from P*livJ* and P*trc** promoters.**

Data analysis and statistics to Figure 2:

Promoter activities determined by real-time measurements. The time window for mRFU calculations is indicated in parentheses.

Conditions: 30 nM Eσ70, 15 nM Plasmid DNA, 0.5 mM NTPs.

Formula to calculate mRFU/min:

mRFU/min = $\frac{(RFUt2-RFUt1)}{(t2-t1)} * 1000$

RFU: relative fluorescence units

t: time after initiation of transcription in min

##### Calculated transcriptional activities (mRFU/min)

**P*livJ***

| *rrnB* terminators | time window (min) | n | mRFU/min Mean | Std. Deviation |
|---|---|---|---|---|
| zero | 20-80 | 4 | 25.52 | 1.66 |
| | 80-140 | 4 | 22.26 | 2.34 |
| | 140-200 | 4 | 20.76 | 2.04 |
| | 200-260 | 4 | 19.00 | 1.78 |
| one | 20-80 | 4 | 20.77 | 0.80 |
| | 80-140 | 4 | 11.84 | 0.77 |
| | 140-200 | 4 | 9.08 | 0.47 |
| | 200-260 | 4 | 7.15 | 0.65 |
| two | 20-80 | 4 | 16.06 | 0.59 |
| | 80-140 | 4 | 8.58 | 0.68 |
| | 140-200 | 4 | 6.38 | 0.63 |
| | 200-260 | 4 | 4.54 | 0.87 |

**P*trc****

| *rrnB* terminators | time window (min) | n | mRFU/min Mean | Std. Deviation |
|---|---|---|---|---|
| zero | 20-80 | 4 | 200.11 | 19.85 |
| | 80-140 | 4 | 158.02 | 22.98 |
| | 140-200 | 4 | 145.03 | 25.31 |
| | 200-260 | 4 | 129.46 | 22.43 |
| one | 20-80 | 4 | 182.68 | 8.19 |
| | 80-140 | 4 | 178.78 | 15.52 |
| | 140-200 | 4 | 179.69 | 17.15 |
| | 200-260 | 4 | 166.46 | 19.53 |
| two | 20-80 | 4 | 187.21 | 10.52 |
| | 80-140 | 4 | 183.14 | 12.37 |
| | 140-200 | 4 | 182.78 | 13.85 |
| | 200-260 | 4 | 173.01 | 11.88 |

Two way ANOVA - Bonferroni Post-hoc-Tests - *rrnB* terminators

**P*livJ***

p-value adjusted for comparison of 12 groups.

| condition 1 | condition 2 | Mean Difference | SE | t | p |
|---|---|---|---|---|---|
| mRFU/min (20-80) - zero | mRFU/min (20-80) - one | 4.75 | 0.90 | 5.29 | <.001 |
| mRFU/min (20-80) - zero | mRFU/min (20-80) - two | 9.46 | 0.90 | 10.53 | <.001 |
| mRFU/min (20-80) - zero | mRFU/min (80-140) - zero | 3.27 | 0.90 | 3.63 | .057 |
| mRFU/min (20-80) - zero | mRFU/min (80-140) - one | 13.69 | 0.90 | 15.23 | <.001 |
| mRFU/min (20-80) - zero | mRFU/min (80-140) - two | 16.95 | 0.90 | 18.86 | <.001 |
| mRFU/min (20-80) - zero | mRFU/min (140-200) - zero | 4.77 | 0.90 | 5.30 | <.001 |
| mRFU/min (20-80) - zero | mRFU/min (140-200) - one | 16.44 | 0.90 | 18.29 | <.001 |
| mRFU/min (20-80) - zero | mRFU/min (140-200) - two | 19.15 | 0.90 | 21.30 | <.001 |

| | | | | | |
|---|---|---|---|---|---|
| mRFU/min (20-80) - zero | mRFU/min (200-260) - zero | 6.53 | 0.90 | 7.26 | <.001 |
| mRFU/min (20-80) - zero | mRFU/min (200-260) - one | 18.37 | 0.90 | 20.44 | <.001 |
| mRFU/min (20-80) - zero | mRFU/min (200-260) - two | 20.99 | 0.90 | 23.35 | <.001 |
| mRFU/min (20-80) - one | mRFU/min (20-80) - two | 4.71 | 0.90 | 5.24 | <.001 |
| mRFU/min (20-80) - one | mRFU/min (80-140) - zero | -1.48 | 0.90 | -1.65 | 1 |
| mRFU/min (20-80) - one | mRFU/min (80-140) - one | 8.94 | 0.90 | 9.94 | <.001 |
| mRFU/min (20-80) - one | mRFU/min (80-140) - two | 12.20 | 0.90 | 13.57 | <.001 |
| mRFU/min (20-80) - one | mRFU/min (140-200) - zero | 0.02 | 0.90 | 0.02 | 1 |
| mRFU/min (20-80) - one | mRFU/min (140-200) - one | 11.69 | 0.90 | 13.01 | <.001 |
| mRFU/min (20-80) - one | mRFU/min (140-200) - two | 14.40 | 0.90 | 16.02 | <.001 |
| mRFU/min (20-80) - one | mRFU/min (200-260) - zero | 1.78 | 0.90 | 1.98 | 1 |
| mRFU/min (20-80) - one | mRFU/min (200-260) - one | 13.62 | 0.90 | 15.16 | <.001 |
| mRFU/min (20-80) - one | mRFU/min (200-260) - two | 16.24 | 0.90 | 18.07 | <.001 |
| mRFU/min (20-80) - two | mRFU/min (80-140) - zero | -6.20 | 0.90 | -6.90 | <.001 |
| mRFU/min (20-80) - two | mRFU/min (80-140) - one | 4.22 | 0.90 | 4.70 | .003 |
| mRFU/min (20-80) - two | mRFU/min (80-140) - two | 7.49 | 0.90 | 8.33 | <.001 |
| mRFU/min (20-80) - two | mRFU/min (140-200) - zero | -4.70 | 0.90 | -5.23 | <.001 |
| mRFU/min (20-80) - two | mRFU/min (140-200) - one | 6.98 | 0.90 | 7.76 | <.001 |
| mRFU/min (20-80) - two | mRFU/min (140-200) - two | 9.68 | 0.90 | 10.77 | <.001 |
| mRFU/min (20-80) - two | mRFU/min (200-260) - zero | -2.94 | 0.90 | -3.27 | .157 |
| mRFU/min (20-80) - two | mRFU/min (200-260) - one | 8.91 | 0.90 | 9.91 | <.001 |
| mRFU/min (20-80) - two | mRFU/min (200-260) - two | 11.53 | 0.90 | 12.82 | <.001 |

| | | | | | |
|---|---|---|---|---|---|
| mRFU/min (80-140) - zero | mRFU/min (80-140) - one | 10.42 | 0.90 | 11.60 | <.001 |
| mRFU/min (80-140) - zero | mRFU/min (80-140) - two | 13.68 | 0.90 | 15.23 | <.001 |
| mRFU/min (80-140) - zero | mRFU/min (140-200) - zero | 1.50 | 0.90 | 1.67 | 1 |
| mRFU/min (80-140) - zero | mRFU/min (140-200) - one | 13.18 | 0.90 | 14.66 | <.001 |
| mRFU/min (80-140) - zero | mRFU/min (140-200) - two | 15.88 | 0.90 | 17.67 | <.001 |
| mRFU/min (80-140) - zero | mRFU/min (200-260) - zero | 3.26 | 0.90 | 3.63 | .058 |
| mRFU/min (80-140) - zero | mRFU/min (200-260) - one | 15.11 | 0.90 | 16.81 | <.001 |
| mRFU/min (80-140) - zero | mRFU/min (200-260) - two | 17.72 | 0.90 | 19.72 | <.001 |
| mRFU/min (80-140) - one | mRFU/min (80-140) - two | 3.26 | 0.90 | 3.63 | .058 |
| mRFU/min (80-140) - one | mRFU/min (140-200) - zero | -8.92 | 0.90 | -9.93 | <.001 |
| mRFU/min (80-140) - one | mRFU/min (140-200) - one | 2.76 | 0.90 | 3.07 | .271 |
| mRFU/min (80-140) - one | mRFU/min (140-200) - two | 5.46 | 0.90 | 6.08 | <.001 |
| mRFU/min (80-140) - one | mRFU/min (200-260) - zero | -7.16 | 0.90 | -7.97 | <.001 |
| mRFU/min (80-140) - one | mRFU/min (200-260) - one | 4.69 | 0.90 | 5.22 | <.001 |
| mRFU/min (80-140) - one | mRFU/min (200-260) - two | 7.30 | 0.90 | 8.13 | <.001 |
| mRFU/min (80-140) - two | mRFU/min (140-200) - zero | -12.18 | 0.90 | -13.56 | <.001 |
| mRFU/min (80-140) - two | mRFU/min (140-200) - one | -0.51 | 0.90 | -0.56 | 1 |
| mRFU/min (80-140) - two | mRFU/min (140-200) - two | 2.20 | 0.90 | 2.45 | 1 |
| mRFU/min (80-140) - two | mRFU/min (200-260) - zero | -10.42 | 0.90 | -11.60 | <.001 |
| mRFU/min (80-140) - two | mRFU/min (200-260) - one | 1.42 | 0.90 | 1.59 | 1 |
| mRFU/min (80-140) - two | mRFU/min (200-260) - two | 4.04 | 0.90 | 4.50 | .004 |
| mRFU/min (140-200) - zero | mRFU/min (140-200) - one | 11.68 | 0.90 | 12.99 | <.001 |

| | | | | | |
|---|---|---|---|---|---|
| mRFU/min (140-200) - zero | mRFU/min (140-200) - two | 14.38 | 0.90 | 16.00 | <.001 |
| mRFU/min (140-200) - zero | mRFU/min (200-260) - zero | 1.76 | 0.90 | 1.96 | 1 |
| mRFU/min (140-200) - zero | mRFU/min (200-260) - one | 13.61 | 0.90 | 15.14 | <.001 |
| mRFU/min (140-200) - zero | mRFU/min (200-260) - two | 16.22 | 0.90 | 18.05 | <.001 |
| mRFU/min (140-200) - one | mRFU/min (140-200) - two | 2.71 | 0.90 | 3.01 | .314 |
| mRFU/min (140-200) - one | mRFU/min (200-260) - zero | -9.92 | 0.90 | -11.03 | <.001 |
| mRFU/min (140-200) - one | mRFU/min (200-260) - one | 1.93 | 0.90 | 2.15 | 1 |
| mRFU/min (140-200) - one | mRFU/min (200-260) - two | 4.55 | 0.90 | 5.06 | .001 |
| mRFU/min (140-200) - two | mRFU/min (200-260) - zero | -12.62 | 0.90 | -14.04 | <.001 |
| mRFU/min (140-200) - two | mRFU/min (200-260) - one | -0.77 | 0.90 | -0.86 | 1 |
| mRFU/min (140-200) - two | mRFU/min (200-260) - two | 1.84 | 0.90 | 2.05 | 1 |
| mRFU/min (200-260) - zero | mRFU/min (200-260) - one | 11.85 | 0.90 | 13.18 | <.001 |
| mRFU/min (200-260) - zero | mRFU/min (200-260) - two | 14.46 | 0.90 | 16.09 | <.001 |
| mRFU/min (200-260) - one | mRFU/min (200-260) - two | 2.62 | 0.90 | 2.91 | .407 |

**P*trc**** 
p-value adjusted for comparison of 12 groups.

| condition 1 | condition 2 | Mean Difference | SE | t | p |
|---|---|---|---|---|---|
| mRFU/min (20-80) - zero | mRFU/min (20-80) - one | 17.43 | 12.33 | 1.41 | 1 |
| mRFU/min (20-80) - zero | mRFU/min (20-80) - two | 12.89 | 12.33 | 1.05 | 1 |
| mRFU/min (20-80) - zero | mRFU/min (80-140) - zero | 42.09 | 12.33 | 3.41 | .106 |
| mRFU/min (20-80) - zero | mRFU/min (80-140) - one | 21.32 | 12.33 | 1.73 | 1 |

| | | | | | |
|---|---|---|---|---|---|
| mRFU/min (20-80) - zero | mRFU/min (80-140) - two | 16.96 | 12.33 | 1.38 | 1 |
| mRFU/min (20-80) - zero | mRFU/min (140-200) - zero | 55.08 | 12.33 | 4.47 | .005 |
| mRFU/min (20-80) - zero | mRFU/min (140-200) - one | 20.41 | 12.33 | 1.66 | 1 |
| mRFU/min (20-80) - zero | mRFU/min (140-200) - two | 17.32 | 12.33 | 1.41 | 1 |
| mRFU/min (20-80) - zero | mRFU/min (200-260) - zero | 70.64 | 12.33 | 5.73 | <.001 |
| mRFU/min (20-80) - zero | mRFU/min (200-260) - one | 33.65 | 12.33 | 2.73 | .643 |
| mRFU/min (20-80) - zero | mRFU/min (200-260) - two | 27.09 | 12.33 | 2.20 | 1 |
| mRFU/min (20-80) - one | mRFU/min (20-80) - two | -4.53 | 12.33 | -0.37 | 1 |
| mRFU/min (20-80) - one | mRFU/min (80-140) - zero | 24.66 | 12.33 | 2.00 | 1 |
| mRFU/min (20-80) - one | mRFU/min (80-140) - one | 3.90 | 12.33 | 0.32 | 1 |
| mRFU/min (20-80) - one | mRFU/min (80-140) - two | -0.47 | 12.33 | -0.04 | 1 |
| mRFU/min (20-80) - one | mRFU/min (140-200) - zero | 37.65 | 12.33 | 3.05 | .279 |
| mRFU/min (20-80) - one | mRFU/min (140-200) - one | 2.99 | 12.33 | 0.24 | 1 |
| mRFU/min (20-80) - one | mRFU/min (140-200) - two | -0.10 | 12.33 | -0.01 | 1 |
| mRFU/min (20-80) - one | mRFU/min (200-260) - zero | 53.22 | 12.33 | 4.32 | .008 |
| mRFU/min (20-80) - one | mRFU/min (200-260) - one | 16.22 | 12.33 | 1.32 | 1 |
| mRFU/min (20-80) - one | mRFU/min (200-260) - two | 9.67 | 12.33 | 0.78 | 1 |
| mRFU/min (20-80) - two | mRFU/min (80-140) - zero | 29.19 | 12.33 | 2.37 | 1 |
| mRFU/min (20-80) - two | mRFU/min (80-140) - one | 8.43 | 12.33 | 0.68 | 1 |
| mRFU/min (20-80) - two | mRFU/min (80-140) - two | 4.07 | 12.33 | 0.33 | 1 |
| mRFU/min (20-80) - two | mRFU/min (140-200) - zero | 42.18 | 12.33 | 3.42 | .103 |
| mRFU/min (20-80) - two | mRFU/min (140-200) - one | 7.52 | 12.33 | 0.61 | 1 |

| | | | | | |
|---|---|---|---|---|---|
| mRFU/min (20-80) - two | mRFU/min (140-200) - two | 4.43 | 12.33 | 0.36 | 1 |
| mRFU/min (20-80) - two | mRFU/min (200-260) - zero | 57.75 | 12.33 | 4.68 | .003 |
| mRFU/min (20-80) - two | mRFU/min (200-260) - one | 20.75 | 12.33 | 1.68 | 1 |
| mRFU/min (20-80) - two | mRFU/min (200-260) - two | 14.20 | 12.33 | 1.15 | 1 |
| mRFU/min (80-140) - zero | mRFU/min (80-140) - one | -20.77 | 12.33 | -1.68 | 1 |
| mRFU/min (80-140) - zero | mRFU/min (80-140) - two | -25.13 | 12.33 | -2.04 | 1 |
| mRFU/min (80-140) - zero | mRFU/min (140-200) - zero | 12.99 | 12.33 | 1.05 | 1 |
| mRFU/min (80-140) - zero | mRFU/min (140-200) - one | -21.68 | 12.33 | -1.76 | 1 |
| mRFU/min (80-140) - zero | mRFU/min (140-200) - two | -24.76 | 12.33 | -2.01 | 1 |
| mRFU/min (80-140) - zero | mRFU/min (200-260) - zero | 28.55 | 12.33 | 2.32 | 1 |
| mRFU/min (80-140) - zero | mRFU/min (200-260) - one | -8.44 | 12.33 | -0.68 | 1 |
| mRFU/min (80-140) - zero | mRFU/min (200-260) - two | -15.00 | 12.33 | -1.22 | 1 |
| mRFU/min (80-140) - one | mRFU/min (80-140) - two | -4.36 | 12.33 | -0.35 | 1 |
| mRFU/min (80-140) - one | mRFU/min (140-200) - zero | 33.75 | 12.33 | 2.74 | .63 |
| mRFU/min (80-140) - one | mRFU/min (140-200) - one | -0.91 | 12.33 | -0.07 | 1 |
| mRFU/min (80-140) - one | mRFU/min (140-200) - two | -4.00 | 12.33 | -0.32 | 1 |
| mRFU/min (80-140) - one | mRFU/min (200-260) - zero | 49.32 | 12.33 | 4.00 | .02 |
| mRFU/min (80-140) - one | mRFU/min (200-260) - one | 12.33 | 12.33 | 1.00 | 1 |
| mRFU/min (80-140) - one | mRFU/min (200-260) - two | 5.77 | 12.33 | 0.47 | 1 |
| mRFU/min (80-140) - two | mRFU/min (140-200) - zero | 38.12 | 12.33 | 3.09 | .252 |
| mRFU/min (80-140) - two | mRFU/min (140-200) - one | 3.45 | 12.33 | 0.28 | 1 |
| mRFU/min (80-140) - two | mRFU/min (140-200) - two | 0.36 | 12.33 | 0.03 | 1 |

| | | | | | |
|---|---|---|---|---|---|
| mRFU/min (80-140) - two | mRFU/min (200-260) - zero | 53.68 | 12.33 | 4.35 | .007 |
| mRFU/min (80-140) - two | mRFU/min (200-260) - one | 16.69 | 12.33 | 1.35 | 1 |
| mRFU/min (80-140) - two | mRFU/min (200-260) - two | 10.13 | 12.33 | 0.82 | 1 |
| mRFU/min (140-200) - zero | mRFU/min (140-200) - one | -34.66 | 12.33 | -2.81 | .523 |
| mRFU/min (140-200) - zero | mRFU/min (140-200) - two | -37.75 | 12.33 | -3.06 | .273 |
| mRFU/min (140-200) - zero | mRFU/min (200-260) - zero | 15.56 | 12.33 | 1.26 | 1 |
| mRFU/min (140-200) - zero | mRFU/min (200-260) - one | -21.43 | 12.33 | -1.74 | 1 |
| mRFU/min (140-200) - zero | mRFU/min (200-260) - two | -27.98 | 12.33 | -2.27 | 1 |
| mRFU/min (140-200) - one | mRFU/min (140-200) - two | -3.09 | 12.33 | -0.25 | 1 |
| mRFU/min (140-200) - one | mRFU/min (200-260) - zero | 50.23 | 12.33 | 4.08 | .016 |
| mRFU/min (140-200) - one | mRFU/min (200-260) - one | 13.24 | 12.33 | 1.07 | 1 |
| mRFU/min (140-200) - one | mRFU/min (200-260) - two | 6.68 | 12.33 | 0.54 | 1 |
| mRFU/min (140-200) - two | mRFU/min (200-260) - zero | 53.32 | 12.33 | 4.33 | .008 |
| mRFU/min (140-200) - two | mRFU/min (200-260) - one | 16.32 | 12.33 | 1.32 | 1 |
| mRFU/min (140-200) - two | mRFU/min (200-260) - two | 9.77 | 12.33 | 0.79 | 1 |
| mRFU/min (200-260) - zero | mRFU/min (200-260) - one | -36.99 | 12.33 | -3.00 | .321 |
| mRFU/min (200-260) - zero | mRFU/min (200-260) - two | -43.55 | 12.33 | -3.53 | .075 |
| mRFU/min (200-260) - one | mRFU/min (200-260) - two | -6.56 | 12.33 | -0.53 | 1 |

#### Data S2 (Fig.4):

**Data analysis and statistics to Figure 4:**

Promoter activities determined by real-time measurements. The time window for mRFU calculations is indicated in parentheses.

Conditions: 30 nM Eσ70, 15 nM Plasmid DNA, 0.5 mM NTPs.

Formula to calculate mRFU/min:

mRFU/min = $\frac{(RFUt2-RFUt1)}{(t2-t1)} * 1000$

RFU: relative fluorescence units
t: time after initiation of transcription in min
"normal" means no Rifampicin added.

##### A: Rifampicin added to the reaction buffer.

**Calculated transcriptional activities (mRFU/min)**

| condition | time window (min) | n | mRFU/min Mean | Std. Deviation |
|---|---|---|---|---|
| normal | 20-80 | 4 | 182.70 | 4.87 |
| | 80-140 | 4 | 177.27 | 2.16 |
| | 140-200 | 4 | 171.84 | 5.95 |
| | 200-260 | 4 | 166.43 | 12.32 |
| 1 µM Rif Buffer | 20-80 | 4 | 1.43 | 0.46 |
| | 80-140 | 4 | 1.39 | 0.24 |
| | 140-200 | 4 | 1.35 | 0.11 |
| | 200-260 | 4 | 1.29 | 0.09 |
| 5 µM Rif Buffer | 20-80 | 4 | 0.85 | 0.17 |
| | 80-140 | 4 | 1.00 | 0.32 |
| | 140-200 | 4 | 1.08 | 0.36 |
| | 200-260 | 4 | 1.08 | 0.09 |

Two way ANOVA - Bonferroni Post-hoc-Tests - Rifampicin concentration

| | | Mean Difference | SE | t | p |
|---|---|---|---|---|---|
| normal | 1 µM Rif Buffer | **173.19** | **1.50** | **115.43** | **<.001** |
| normal | 5 µM Rif Buffer | **173.56** | **1.50** | **115.67** | **<.001** |
| 1 µM Rif Buffer | 5 µM Rif Buffer | **0.36** | **1.50** | **0.24** | **1** |

**p-value adjusted for comparison of 3 groups.**

#### B: Rifampicin added to NTPs.

##### Calculated transcriptional activities (mRFU/min)

| condition | time window | n | mRFU/min Mean | Std. Deviation |
|---|---|---|---|---|
| normal | 20-80 | 4 | 172.81 | 29.03 |
| | 80-140 | 4 | 152.83 | 29.72 |
| | 140-200 | 4 | 139.17 | 30.00 |
| | 200-260 | 4 | 131.82 | 29.92 |
| 1 µM Rif NTPs | 20-80 | 4 | 14.43 | 1.20 |
| | 80-140 | 4 | 6.04 | 0.59 |
| | 140-200 | 4 | 2.35 | 0.37 |
| | 200-260 | 4 | 3.35 | 0.47 |
| 5 µM Rif NTPs | 20-80 | 4 | 6.90 | 0.64 |
| | 80-140 | 4 | 3.31 | 0.56 |
| | 140-200 | 4 | 1.54 | 0.39 |
| | 200-260 | 4 | 1.60 | 0.16 |

Two way ANOVA - Bonferroni Post-hoc-Tests - Rifampicin concentration

| | | Mean Difference | SE | t | p |
|---|---|---|---|---|---|
| normal | 1 µM Rif NTPs | 142.61 | 6.06 | 23.54 | <.001 |
| normal | 5 µM Rif NTPs | 145.82 | 6.06 | 24.07 | <.001 |
| 1 µM Rif NTPs | 5 µM Rif NTPs | 3.20 | 6.06 | 0.53 | 1 |

p-value adjusted for comparison of 3 groups.

#### C: Rifampicin added during IVT at 123 min.

##### Calculated transcriptional activities (mRFU/min)

| condition | time window | n | mRFU/min Mean | Std. Deviation |
|---|---|---|---|---|
| normal | 20-80 | 4 | 174.65 | 18.14 |
| | 140-200 | 4 | 103.02 | 15.74 |
| | 200-260 | 4 | 90.82 | 14.80 |
| 1 µM Rif at 120 min | 20-80 | 4 | 182.10 | 11.22 |
| | 140-200 | 4 | 24.17 | 4.87 |
| | 200-260 | 4 | 12.73 | 1.22 |
| 5 µM Rif at 120 min | 20-80 | 4 | 170.69 | 9.33 |
| | 140-200 | 4 | 20.17 | 3.64 |
| | 200-260 | 4 | 9.68 | 1.13 |

Two way ANOVA - Bonferroni Post-hoc-Tests

| condition 1 | condition 2 | Mean Difference | SE | t | p |
|---|---|---|---|---|---|
| mRFU/min 20-80 - normal | mRFU/min 20-80 - 1 µM Rif at 120 min | -7.45 | 7.63 | -0.98 | 1 |
| mRFU/min 20-80 - normal | mRFU/min 20-80 - 5 µM Rif at 120 min | 3.96 | 7.63 | 0.52 | 1 |

| | | | | | |
|---|---|---|---|---|---|
| mRFU/min 20-80 - normal | mRFU/min 140-200 - normal | 71.63 | 7.63 | 9.39 | <.001 |
| mRFU/min 20-80 - normal | mRFU/min 140-200 - 1 µM Rif at 120 min | 150.48 | 7.63 | 19.72 | <.001 |
| mRFU/min 20-80 - normal | mRFU/min 140-200 - 5 µM Rif at 120 min | 154.48 | 7.63 | 20.24 | <.001 |
| mRFU/min 20-80 - normal | mRFU/min 200-260 - normal | 83.83 | 7.63 | 10.98 | <.001 |
| mRFU/min 20-80 - normal | mRFU/min 200-260 - 1 µM Rif at 120 min | 161.92 | 7.63 | 21.22 | <.001 |
| mRFU/min 20-80 - normal | mRFU/min 200-260 - 5 µM Rif at 120 min | 164.97 | 7.63 | 21.62 | <.001 |
| mRFU/min 20-80 - 1 µM Rif at 120 min | mRFU/min 20-80 - 5 µM Rif at 120 min | 11.41 | 7.63 | 1.50 | 1 |
| mRFU/min 20-80 - 1 µM Rif at 120 min | mRFU/min 140-200 - normal | 79.08 | 7.63 | 10.36 | <.001 |
| mRFU/min 20-80 - 1 µM Rif at 120 min | mRFU/min 140-200 - 1 µM Rif at 120 min | 157.93 | 7.63 | 20.69 | <.001 |
| mRFU/min 20-80 - 1 µM Rif at 120 min | mRFU/min 140-200 - 5 µM Rif at 120 min | 161.93 | 7.63 | 21.22 | <.001 |
| mRFU/min 20-80 - 1 µM Rif at 120 min | mRFU/min 200-260 - normal | 91.29 | 7.63 | 11.96 | <.001 |
| mRFU/min 20-80 - 1 µM Rif at 120 min | mRFU/min 200-260 - 1 µM Rif at 120 min | 169.38 | 7.63 | 22.19 | <.001 |
| mRFU/min 20-80 - 1 µM Rif at 120 min | mRFU/min 200-260 - 5 µM Rif at 120 min | 172.43 | 7.63 | 22.59 | <.001 |
| mRFU/min 20-80 - 5 µM Rif at 120 min | mRFU/min 140-200 - normal | 67.67 | 7.63 | 8.87 | <.001 |
| mRFU/min 20-80 - 5 µM Rif at 120 min | mRFU/min 140-200 - 1 µM Rif at 120 min | 146.52 | 7.63 | 19.20 | <.001 |
| mRFU/min 20-80 - 5 µM Rif at 120 min | mRFU/min 140-200 - 5 µM Rif at 120 min | 150.52 | 7.63 | 19.72 | <.001 |
| mRFU/min 20-80 - 5 µM Rif at 120 min | mRFU/min 200-260 - normal | 79.88 | 7.63 | 10.47 | <.001 |
| mRFU/min 20-80 - 5 µM Rif at 120 min | mRFU/min 200-260 - 1 µM Rif at 120 min | 157.97 | 7.63 | 20.70 | <.001 |
| mRFU/min 20-80 - 5 µM Rif at 120 min | mRFU/min 200-260 - 5 µM Rif at 120 min | 161.02 | 7.63 | 21.10 | <.001 |
| mRFU/min 140-200 - normal | mRFU/min 140-200 - 1 µM Rif at 120 min | 78.85 | 7.63 | 10.33 | <.001 |
| mRFU/min 140-200 - normal | mRFU/min 140-200 - 5 µM Rif at 120 min | 82.85 | 7.63 | 10.86 | <.001 |
| mRFU/min 140-200 - normal | mRFU/min 200-260 - normal | 12.20 | 7.63 | 1.60 | 1 |

| | | | | | |
|---|---|---|---|---|---|
| mRFU/min 140-200 - normal | mRFU/min 200-260 - 1 µM Rif at 120 min | 90.30 | 7.63 | 11.83 | <.001 |
| mRFU/min 140-200 - normal | mRFU/min 200-260 - 5 µM Rif at 120 min | 93.35 | 7.63 | 12.23 | <.001 |
| mRFU/min 140-200 - 1 µM Rif at 120 min | mRFU/min 140-200 - 5 µM Rif at 120 min | 4.00 | 7.63 | 0.52 | 1 |
| mRFU/min 140-200 - 1 µM Rif at 120 min | mRFU/min 200-260 - normal | -66.64 | 7.63 | -8.73 | <.001 |
| mRFU/min 140-200 - 1 µM Rif at 120 min | mRFU/min 200-260 - 1 µM Rif at 120 min | 11.45 | 7.63 | 1.50 | 1 |
| mRFU/min 140-200 - 1 µM Rif at 120 min | mRFU/min 200-260 - 5 µM Rif at 120 min | 14.50 | 7.63 | 1.90 | 1 |
| mRFU/min 140-200 - 5 µM Rif at 120 min | mRFU/min 200-260 - normal | -70.64 | 7.63 | -9.26 | <.001 |
| mRFU/min 140-200 - 5 µM Rif at 120 min | mRFU/min 200-260 - 1 µM Rif at 120 min | 7.45 | 7.63 | 0.98 | 1 |
| mRFU/min 140-200 - 5 µM Rif at 120 min | mRFU/min 200-260 - 5 µM Rif at 120 min | 10.50 | 7.63 | 1.38 | 1 |
| mRFU/min 200-260 - normal | mRFU/min 200-260 - 1 µM Rif at 120 min | 78.09 | 7.63 | 10.23 | <.001 |
| mRFU/min 200-260 - normal | mRFU/min 200-260 - 5 µM Rif at 120 min | 81.14 | 7.63 | 10.63 | <.001 |
| mRFU/min 200-260 - 1 µM Rif at 120 min | mRFU/min 200-260 - 5 µM Rif at 120 min | 3.05 | 7.63 | 0.40 | 1 |

p-value adjusted for comparison of 9 groups.
